# Systematic identification of functional SNPs interrupting 3’ UTR polyadenylation signals

**DOI:** 10.1101/782425

**Authors:** Eldad David Shulman, Ran Elkon

## Abstract

Alternative polyadenylation (APA) is emerging as a widespread regulatory layer as the majority of human protein-coding genes contain several polyadenylation (p(A)) sites in their 3’ UTRs. By generating isoforms with different 3’ UTR length, APA potentially affects mRNA stability, translation efficiency, nuclear export, and cellular localization. Polyadenylation sites are regulated by adjacent RNA cis-regulatory elements, the principals among them are the polyadenylation signal (PAS) AAUAAA and its main variant AUUAAA, typically located ~20- nt upstream of the p(A) site. Mutations in PAS and other auxiliary poly(A) cis-elements in the 3’ UTR of several genes have been shown to cause human Mendelian diseases, and to date, only a few common SNPs that regulate APA were associated with complex diseases. Here, we systematically searched for SNPs that affect gene expression and human traits by modulation of 3’ UTR APA. Focusing on the variants most likely to exert the strongest effect, we identified 2,305 SNPs that interrupt the canonical PAS or its main variant. Implementing pA-QTL tests using GTEx RNA-seq data, we identified 139 PAS SNPs significantly associated with the usage of their p(A) site. As expected, PAS-interrupting alleles were significantly linked with decreased cleavage at their p(A) site and the consequential 3’ UTR lengthening. As an indication for a functional effect of these PAS SNPs on gene expression, 65 of the pA-QTLs were also detected as eQTLs of the same gene in the same tissue. Furthermore, we observed that PAS-interrupting alleles linked with 3’ UTR lengthening were also strongly associated with decreased gene expression, pointing that shorter isoforms generated by APA are generally more stable than longer ones. Last, indicative of the impact of PAS SNPs on human phenotypes, 53 pA-QTLs overlapped GWAS SNPs that are significantly linked with human traits.

## Introduction

The maturation of mRNA 3′ ends is a 2-steps process, termed *cleavage and polyadenylation*, that involves endonucleolytic cleavage of the nascent RNA followed by synthesis of a poly(A) tail on the 3′ terminus of the cleaved product [1]. Cleavage and polyadenylation sites (p(A) sites) are determined and controlled by adjacent RNA cis-regulatory elements, the principal among them is the *polyadenylation signal* (*PAS*) AAUAAA, typically located ~20-nt upstream of the p(A) site. There are more than 10 weaker variants to this canonical PAS, the main among them is AUUAAA [2]. Auxiliary elements include upstream U-rich and UGUA motifs and downstream U-rich and GU-rich elements, and the strength of a p(A) site is determined by these elements in a combinatorial manner [3]. Importantly, the majority of human protein-coding genes contain several alternative p(A) sites in their 3’ UTR, making alternative polyadenylation (APA) a widespread regulatory layer that generates transcript isoforms with alternative 3′ ends, and correspondingly, different 3’ UTR lengths [1, 4–6]. As 3′ UTRs contain cis-elements that serve as major docking platforms for microRNAs (miRNAs) and RNA binding proteins (RBPs), which are involved in various aspects of mRNA metabolism, 3′ UTR APA can affect post-transcriptional regulation in multiple ways, including the modulation of mRNA stability, translation efficiency, nuclear export and cellular localization [7–9]. Transcriptomic studies demonstrated that APA is globally modulated during multiple differentiation processes [4, 10–17] and in response to changes in cell proliferation state [18–20]. Yet, our current understanding of the impact of APA on gene regulation and of its biological roles is still very rudimentary.

Mutations in PAS and other poly(A) cis-elements, and the resulting alteration in gene expression have been shown to cause several human Mendelian diseases. Examples include the mutation in the 3’ UTR of *HBA2* (converting AATAAA to AATAAG) causing α-Thalassaemia [21], the mutation in the 3’ UTR of *HBB* (AATAAA to AACAAA) causing β-Thalassaemia [22] and the mutation in the 3’ UTR of *FOXP3* (AATAAA to AATGAA) causing the IPEX syndrome [23] (for a thorough review see [24]). In addition, few common SNPs that regulate APA were found to affect the risk of complex diseases. This includes a risk SNP for systemic lupus erythematosus (SLE) located in the 3’ UTR of *IRF5*. This SNP reduces the use of a proximal p(A) site, leading to the production of longer and less stable isoforms and consequently to reduced *IRF5* levels [25]. Similarly, a polymorphic cis-element downstream to a PAS in the 3’ UTR of *ATP1B1* is associated with high blood pressure [26].

Given the high prevalence of APA, there are likely many more common SNPs that modulate APA and affect disease susceptibility. In this study, we systematically searched for SNPs that interfere with 3’ UTR PAS signals in the human genome and implemented polyadenylation-QTL (pA-QTL) tests to examine their effect on APA. Analyzing GTEx RNA-seq data from twelve tissues we identified dozens of such SNPs that significantly affect the usage of their downstream p(A) sites. The intersection of these pA-QTLs with eQTLs and GWAS risk SNPs indicated their roles in the regulation of gene expression and their impact on various phenotypes.

## Results

We first sought to systematically identify SNPs in the human genome that affect 3’ UTR APA. To focus our analysis on the ones likely to exert the strongest effect on gene regulation (and thus on phenotype), we specifically searched for SNPs that interrupt the canonical PAS (AATAAA) or its main variant (ATTAAA). Using p(A) sites annotations from polyA DB [27], we overall detected 2,305 SNPs that interfere with these signals in a 40-nt window upstream of annotated 3’ UTR p(A) sites in 1,936 unique genes (Figure 1A; **Supplementary Table 1**). We call such variants *PAS SNPs*. Each PAS SNP has one allele that preserves the PAS and another allele that interrupts it (called PAS Interrupting Allele (*PIA*)). Note that the PIA can be either the allele appearing on the genome reference sequence (the reference allele) or the alternative allele (Figure 1B). 68% of the PAS SNPs we detected were located within the canonical AATAAA PAS (**Supplementary Table 1**). We associated each PAS SNP with its downstream p(A) site.

**Figure 1.**
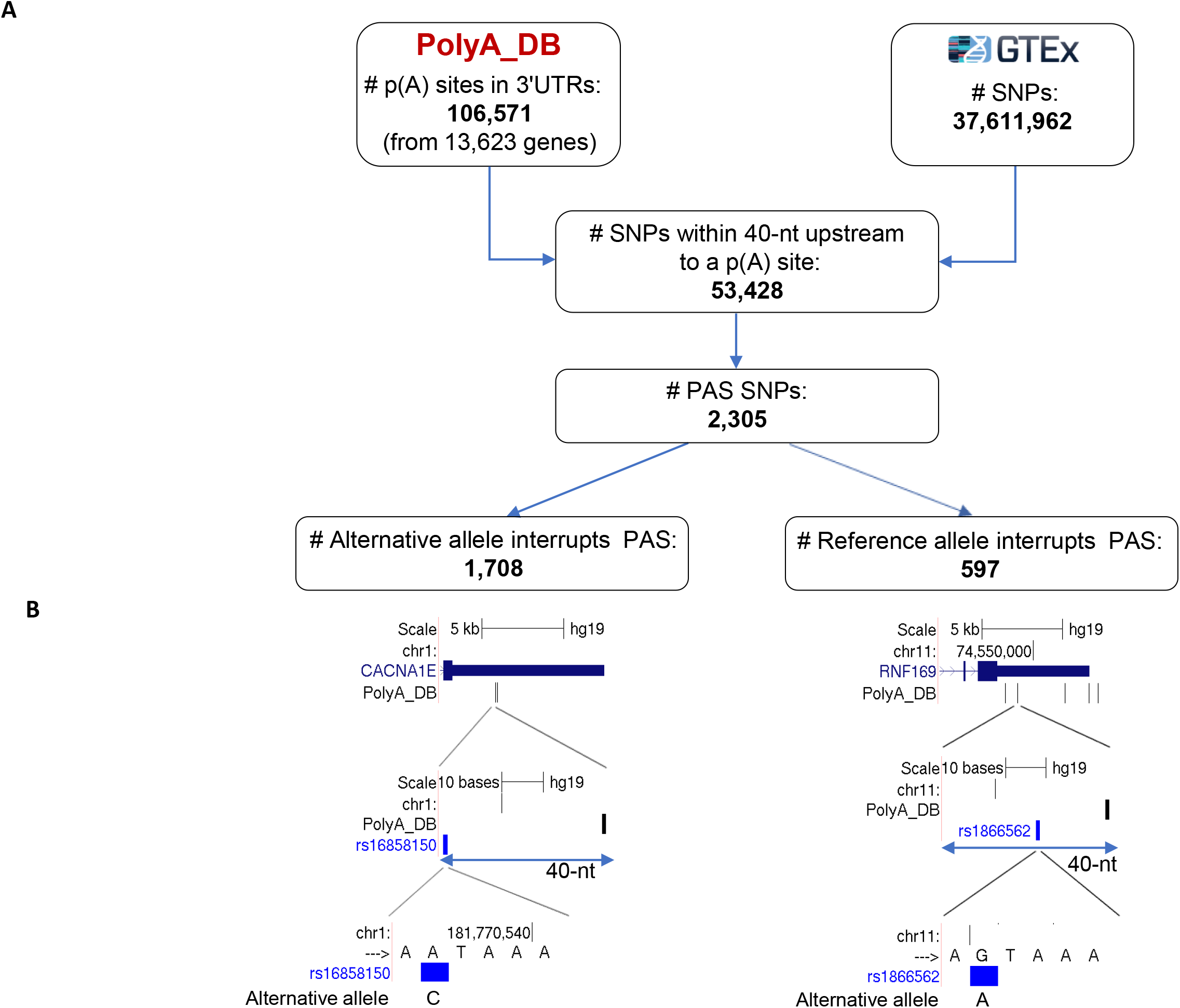
Systematic identification of PAS SNPs in the human genome. **A.** We defined as *PAS SNPs* those that are located within 40-nt upstream of an annotated 3’ UTR p(A) site and have an allele that interrupts the canonical PAS sequence AATAAA or its main variant ATTAAA. We considered all the 3’ UTR p(A) sites annotated in poly(A) DB (release 3.2), and all 37,611,962 SNPs included in GTEX v7. We detected 2,305 such SNPs. Each biallelic SNP has a reference allele (the allele that appears in the genome’s reference sequence) and an alternative allele. Among the 2,305 PAS SNPs detected by our screen, 1,708 SNPs have the alternative allele interrupting the PAS sequence and 597 SNPs have the reference allele disrupting the signal. **B**. An example for a PAS SNP whose alternative allele interrupts the PAS signal (rs16858150 in the 3’ UTR of *CACNA1E*; left) and a PAS SNP whose reference allele interrupts it (rs1866562 in the 3’ UTR of *RNF169*; right).

PAS SNPs potentially have a strong impact on cleavage and polyadenylation efficiency at their downstream p(A) sites. To detect such effects we implemented pA-QTL tests that are based on the estimation of p(A)-site usage from RNA-seq data. Each PAS SNP divides its 3’ UTR into two segments: a common UTR (cUTR) which is upstream to the associated p(A) site and is common to the transcript generated by cleavage at this p(A) site and the transcripts generated by usage of more distal p(A) sites, and an alternative UTR (aUTR) which is included only in transcripts generated by more distal p(A) sites (Figure 2A). For each RNA-seq sample, we estimate the usage of each p(A) site associated with a PAS SNP by calculating the p(A) site Usage Index (*pAUI*), defined as the ratio (in log scale) between the number of reads mapping to the cUTR and the aUTR segments (Figure 2A). Having RNA-seq data from a large cohort of individuals for which genotype data is available too, enables to test for association between the PAS SNP alleles and the pAUI levels (Figure 2A). We carried out such pA-QTL tests on twelve selected tissue from the GTEx project (v7) [28] that have more than 130 RNA-seq samples with corresponding genotype data (Methods). We carried out the pA-QTL analyses on each tissue separately. To ensure sufficient statistical power while lowering the burden of multiple testing, for each tissue, we first filtered out PAS SNPs whose minor allele frequency (MAF) was below 0.01 among the subjects considered for that tissue. In total, 558 PAS SNPs (out of 2,305) passed this criterion, and 139 of them were significantly (FDR=5%) associated with the usage of their p(A) sites. 75% of these 139 pA-QTLs interrupt a canonical PAS (and the rest interrupt the ATTAAA variant) (**Supplementary Table 2-3**). Most of these pA-QTLs were detected in multiple tissues (Figure 2B-D, Figure 3 and **Supplementary Table 2-3**).

**Figure 2.**
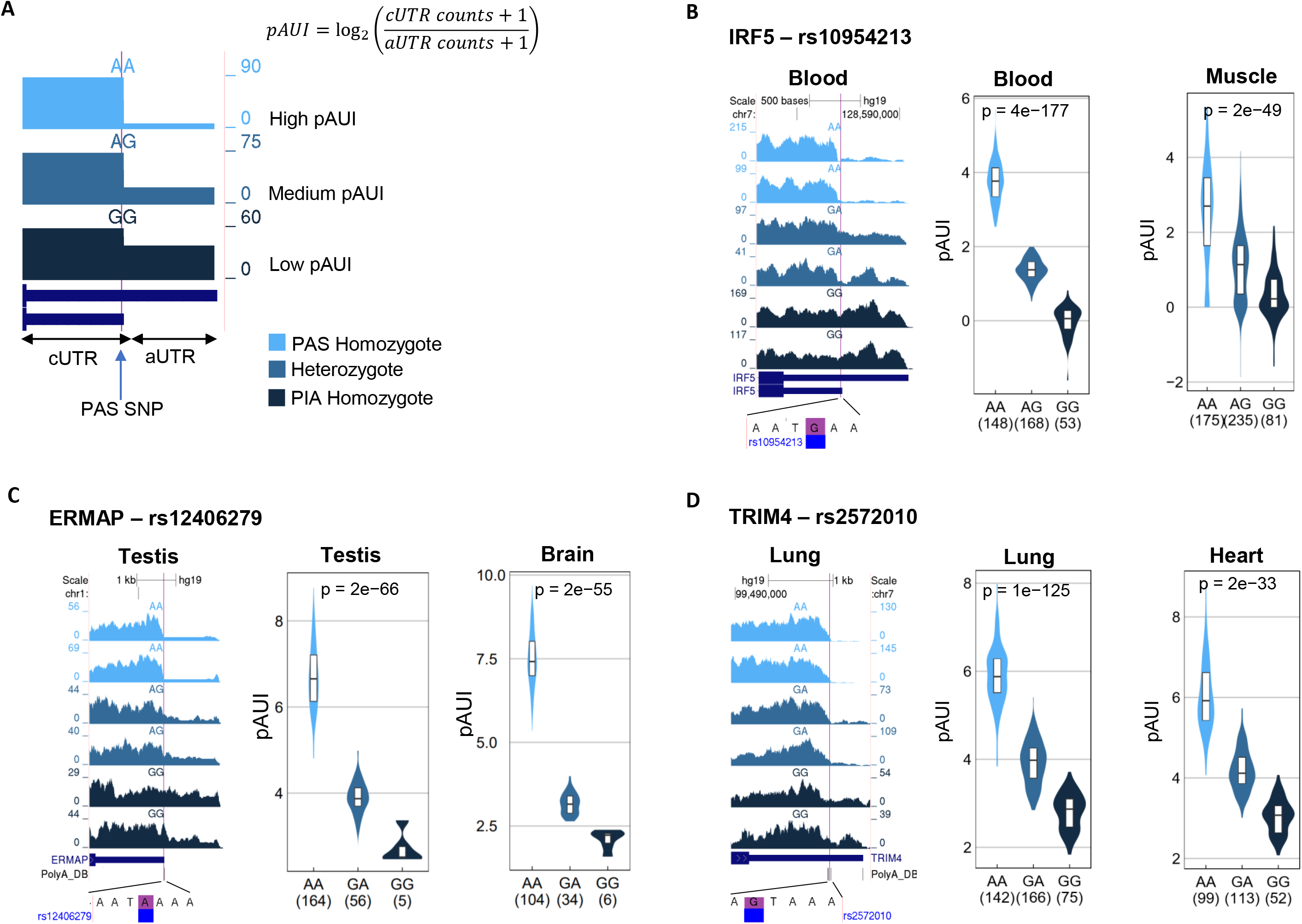
Identification of PAS SNPs that are pA-QTLs. **A**. We used the p(A) site usage index (pAUI) to quantify cleavage efficiency at each annotated 3’ UTR pA site in each RNA-seq sample. The pAUI is defined as the ratio (in log2 scale) between the counts of 3’ UTR reads mapping upstream of the pA site (common 3’ UTR segment; cUTR) and those mapping downstream of it (alternative 3’ UTR segment; aUTR) (Methods). We then used this index to detect PAS SNPs that show a significant association between alleles and pAUI levels of their p(A) site. SNPs showing such association are referred to as pA-QTLs. The expected pattern, as shown in this cartoon, is that the PAS-preserving allele is associated with higher usage of the p(A) site while the PAS-interrupting allele (*PIA*) is associated with reduced usage this site (resulting in 3’ UTR lengthening). Heterozygotes for such SNPs are expected to show intermediate pAUI levels compared to the two homozygotes. The cartoon illustrates reads coverage on a 3’ UTR for three RNA-seq samples of varying levels of pAUI, colored according to the genotype of the PAS SNP. **B**–**D.** Examples of three PAS SNPs consistently detected as pA-QTLs in multiple tissues. In each example, the left panel shows reads coverage in the gene’s 3’ UTR from RNA-seq samples of two selected donors from each PAS SNP genotype. The vertical purple line marks the location of the PAS SNP. The genome reference sequence around the PAS SNP is shown below. Violin plots in the middle and left panels show the distribution of pAUI levels in each genotype group for a given tissue (In each plot, homozygotes of the PAS-preserving allele are shown in the left, heterozygotes – in the middle, and homozygotes to the PAS-interrupting allele (PIA) – in the right violin. The number of individuals in each group is indicated in parentheses). P-values are calculated using linear regression (see Methods). (In C, “Brain” refers to “Brain caudal nucleus”, and in D, “Heart” refers to “Heart atrial appendage”).

**Figure 3.**
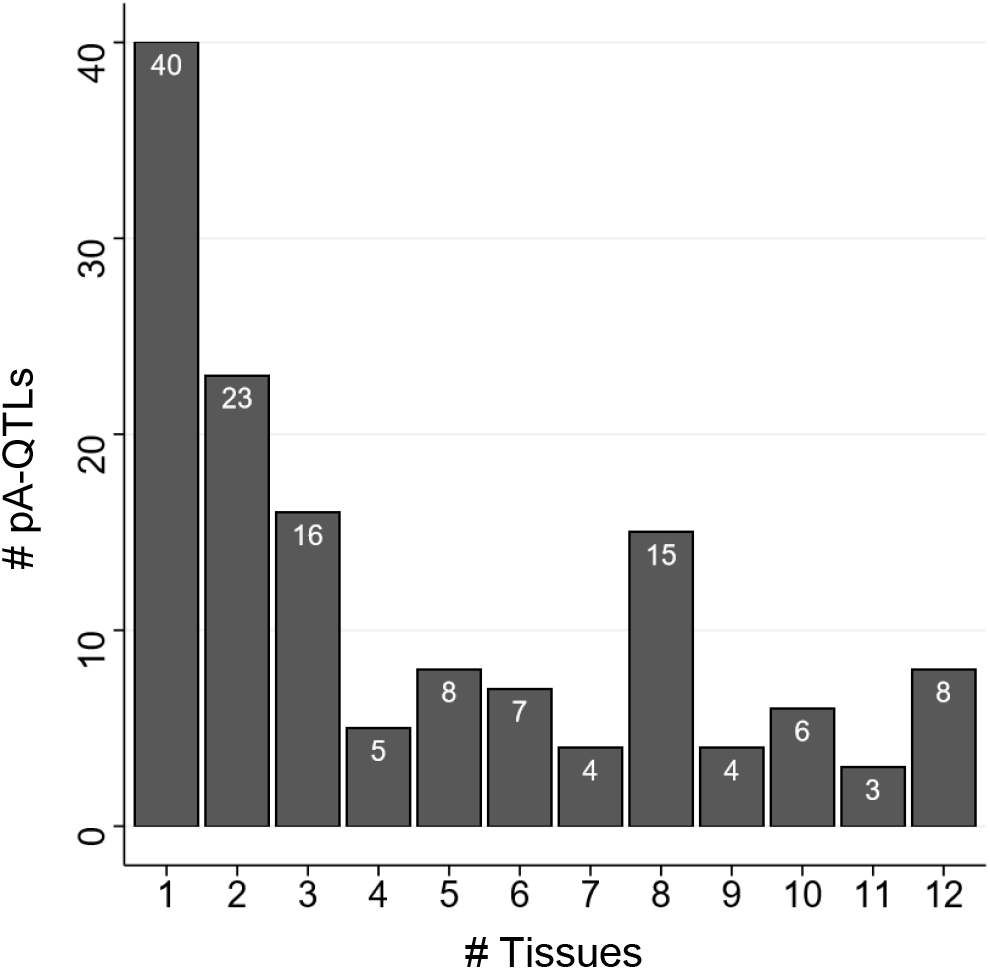
Consistent pA-QTL effects detected in multiple tissues. Distribution of the number of different tissues in which the pA-QTLs were detected. 40 pA-QTLs were detected in a single tissue; 17 pA-QTLs were detected in at least 10 tissues.

As the PAS-interrupting allele (PIA) of a PAS SNP is expected to reduce cleavage efficiency at its p(A) site, we anticipated that the pA-QTL PIAs will be associated with lower pAUI (reflecting 3’ UTR lengthening) (Figure 2A). However, unexpectedly, we also identified pA-QTLs whose PIA was associated with increased pAUI (Figure 4A-B). Interestingly, in these cases, disrupting a p(A) site by the PIA of a PAS SNP caused an increased usage of an upstream (proximal) p(A) site (resulting in an increased pAUI and 3’ UTR shortening). Yet, overall, the cases in which the PAI showed the expected 3’ UTR lengthening effect were markedly more prevalent (88%; Figure 4C). Furthermore, remarkably, for pA-QTLs that were detected in multiple tissues, the effect of their PIA on 3’ UTR length was completely consistent across all the tissues (Figure 4D).

**Figure 4.**
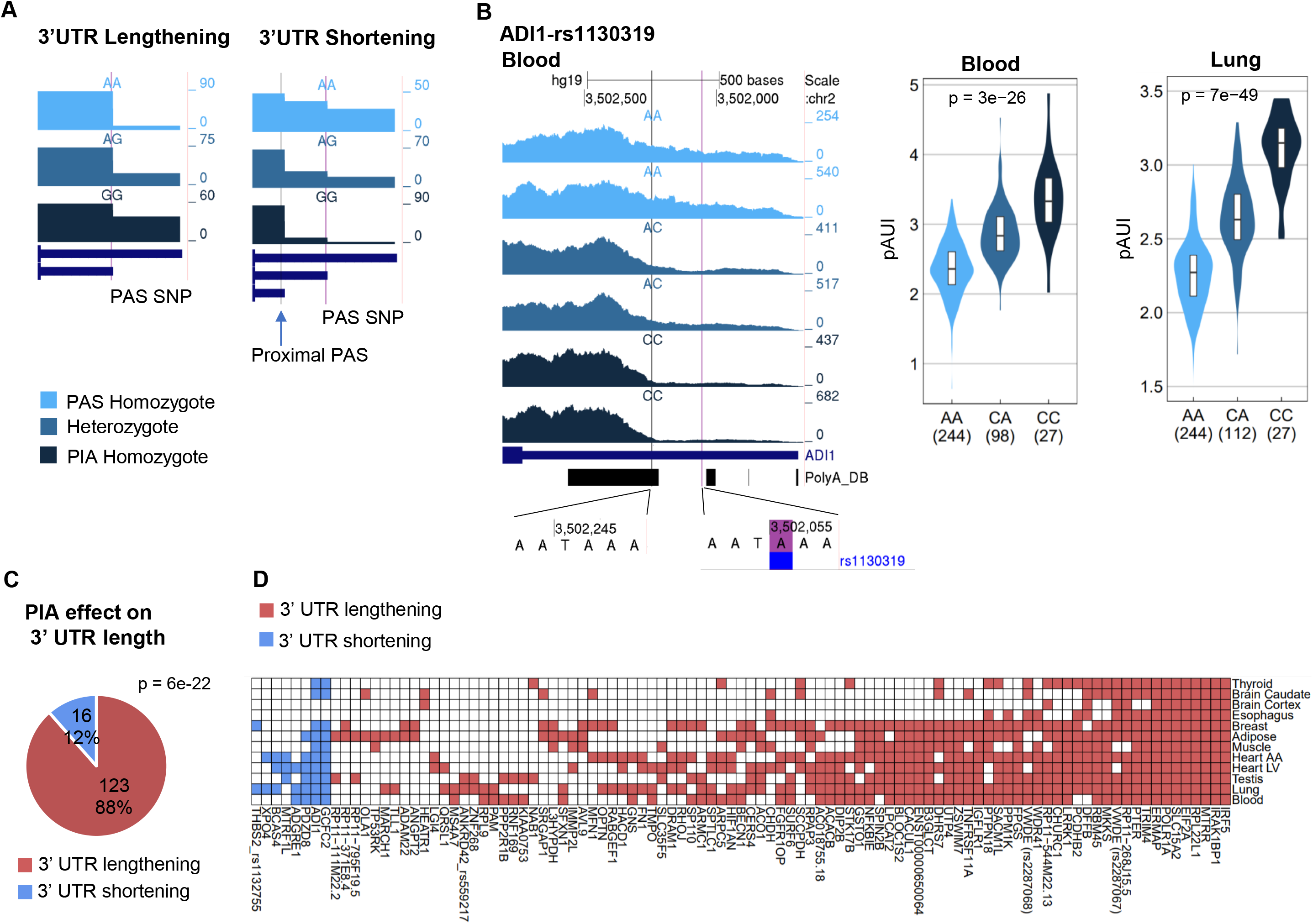
The effect of PAS interrupting alleles (PIAs) on 3’ UTR length. **A**. Cartoons illustrating the anticipated lengthening effect of PIAs on the 3’ UTR (left) and the unexpected 3’ UTR shortening effect, due to elevated usage of an alternative proximal p(A) site (right). Note that in the lengthening case the PIA is associated with decreased pAUI levels whereas in the shortening case, the PIA is associated with elevated pAUI levels. **B**. An example of a PAS SNP (rs1130319 in the 3’ UTR of *ADI1*) whose PIA is associated with 3’ UTR shortening (increased pAUI). Notably, this PAS SNP is detected as a pA-QTL in ten out of the twelve analyzed tissues and in all of these ten tissues, its PIA is consistently associated with 3’ UTR shortening effect (shown in **D**). **C**. A pie chart for the effect of pA-QTL’s PIA on 3’ UTR length. As expected, in the majority of cases the PAI showed a lengthening effect (p-value calculated using single-tailed binomial test). Yet, in 12% of the cases, disrupting the PAS resulted in an increased usage of an upstream 3’ UTR p(A) site (resulting in 3’ UTR shortening). (Note, each pA-QTL is counted once in the pie chart even if it was detected as pA-QTL in multiple tissues.) **D.** Remarkably, PIAs showed a completely consistent effect over all the tissues in which the PAS SNP was detected as a pA-QTL (shown are all PAS SNPs detected as pA-QTL in more than one tissue).

Next, to test for possible functional effects of PAS SNPs on gene expression (e.g., through regulation of transcript stability), we intersected our detected pA-QTLs with GTEx eQTLs (Figure 5A). On average, ~40% (25%-59%) of the pA-QTLs were also detected as eQTLs of the same gene in the same tissue (Figure 5B). Overall, of the 139 pA-QTLs detected by our analyses, 65 were also detected as eQTLs (of the same gene) in at least one common tissue (Figure 5C). Notably, the PAS SNP in the 3’ UTR of *EIF2A* was detected as both pA- QTL and eQTL in all twelve tissues, and the PAS SNP in the 3’ UTR of *TRIM4* was detected as both pA-QTL and eQTL in eleven tissues (Figure 5C).

**Figure 5.**
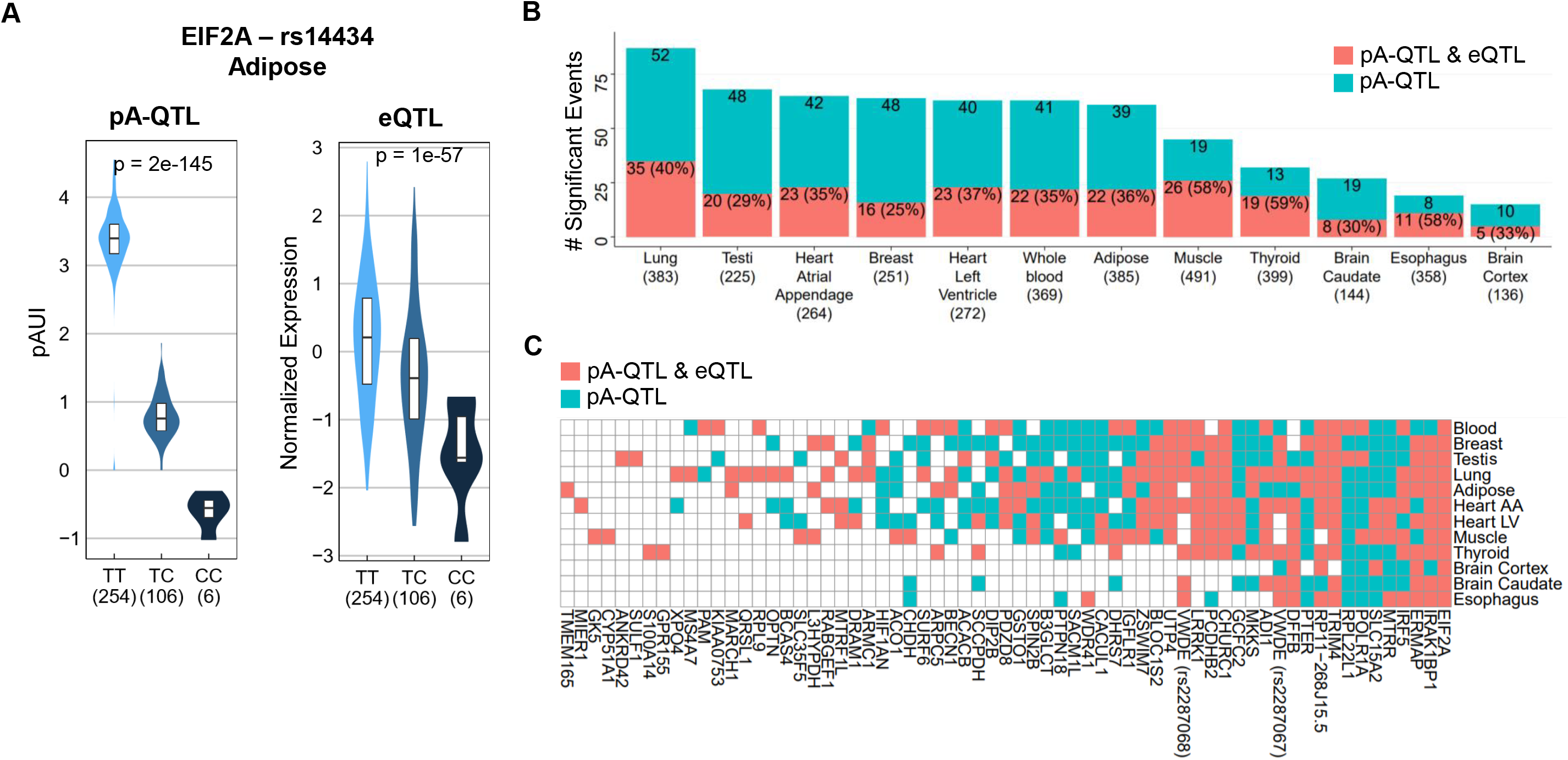
Association between pA-QTLs and eQTLs. **A**. As an example, rs14434 in the 3’ UTR of *EIF2A* is both a pA-QTL and an eQTL of this gene. The PIA of rs14434 (which is the C allele) is associated with both lower pAUI at the corresponding p(A) site (and thus, 3’ UTR lengthening) and lower expression level of *EIF2A* (p-value calculated using linear regression see Methods). Notably, rs14434 showed this same effect in all 12 tissues (Figure 6A). **B.** Bar plot of the number of pA-QTLs detected in each tissue, and the portion of them that are also detected as eQTLs (of the same gene). The number of samples analyzed in each tissue is indicated in parentheses. **C**. The plot shows the 65 PAS SNPs that were detected in at least one tissue as both pA-QTL and eQTL (of the same gene). Each column represents a pA-QTL, labeled by its gene’s symbol, and indicates in which tissues the PAS SNP was detected either as a pA- QTL or as both pA-QTL and eQTL. For example, the PAS SNP in the 3’ UTR of *EIF2A* was detected as both pA-QTL and eQTL in all 12 tissues; the PAS SNP in the 3’ UTR of *ADI1* was detected as pA-QTL in 10 tissues, in 7 of them it was detected also as an eQTL.

We next examined a possible association between the effect of the PIA on 3’ UTR length (lengthening or shortening) and its effect on gene expression (increased or decreased expression). As 3’ UTR cis-regulatory elements mostly have destabilizing effects (e.g., microRNA binding sites), we expected that PIAs associated with 3’ UTR lengthening will consequently be mainly associated with decreased expression level (see Figure 5A for one example). Indeed, we observed that 3’ UTR lengthening effect of the PIAs was significantly linked with reduced expression in eleven (out of twelve) tissues (Figure 6A). Yet, we detected few uncommon cases in which 3’ UTR lengthening effect of a PIA was associated with increased expression level (Figure 6B), and such effects were too consistently observed across different tissues (Figure 6A). On the other hand, 3’ UTR shortening was generally linked with increased gene expression but was not frequent enough to allow robust statistical testing of this tendency (Figure 6A).

**Figure 6.**
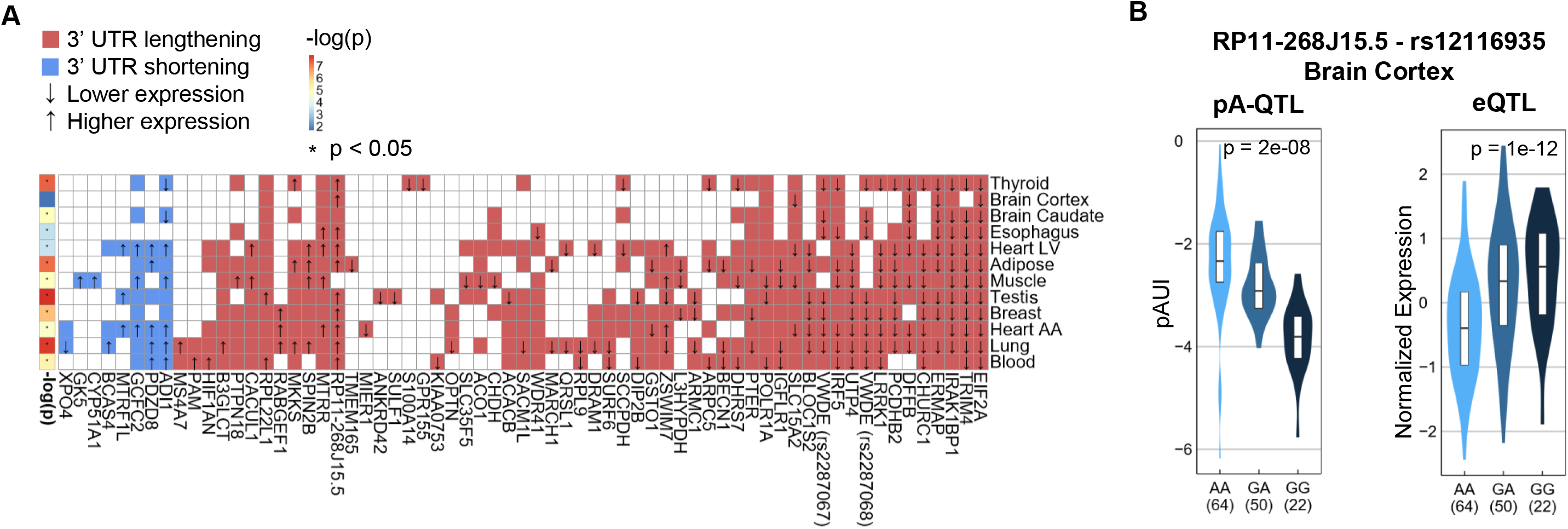
Association between PIA effects on 3’ UTR length and gene expression level. **A.** Heatmap for the 65 pA-QTLs shown in Figure 5C presenting here, for each, the PIA’s effect on 3’ UTR length (color) and, if the PAS SNP was also detected as eQTL, the PIA’s effect on gene expression level (arrow). The left-most column shows, per tissue, the significance of the association between 3’ UTR lengthening and decreased expression level (p-value calculated using one-tailed binomial tests). (There were too few cases of 3’ UTR shortening effect to carry out similar test for the link between 3’ UTR shortening and increased expression, though such general tendency is apparent). **B.** An example of the uncommon case where a PIA (the G allele) is associated with decreased pAUI of the corresponding p(A) site (that is, 3’ UTR lengthening) and with higher expression of the target gene. This PIA shows the same effect in 10 different tissues (**A**).

Last, we examined possible links between our identified pA-QTLs and human phenotypes by intersecting them with GWAS risk SNPs (Methods). In total, 53 (out of 139) pA- QTLs overlapped a GWAS SNP significantly associated with a human trait (**Supplementary Table 3**). This analysis captured the known effect of the IRF5’s PAS SNP on SLE susceptibility [25], in addition to suggesting few novel links. These include possible links between the pA- QTL in the 3’ UTR of *ERMAP* (erythroblast membrane-associated protein) and red cell distribution width (RDW); of the pA-QTLs in the 3’ UTRs of *LPCAT2* (lysophosphatidylcholine acyltransferase 2) and *CERS4* (ceramide synthase 4) and thyroid-stimulating hormone levels; and of the pA-QTL in the 3’ UTR of *ARMC1* (armadillo repeat containing 1) and cognitive performance. Interestingly, the expression pattern of these genes corresponds to the traits with which they are associated: *IRF5* and *ERMAP* show markedly high expression in blood cells; *LPCAT2* and *CERS4* in the thyroid gland and *ARMC1*is most highly expressed in the brain.

## Discussion

Despite the emergence of APA as an important layer of gene regulation, potentially affecting the majority of human protein-coding genes, it remains largely unexplored, and its involvement in physiological and pathological processes is often overlooked. Here, we searched for SNPs that modulate APA. Focusing on SNPs likely to have the strongest effect, we identified 2,305 SNPs that interfere with the canonical PAS or its main variant. Seeking support for the regulatory impact of these SNPs on APA, we implemented pA-QTL tests using GTEx RNA-seq data from twelve tissues. 558 PAS SNPs had MAF level high enough to allow robust statistical examination (MAF>0.1), and 139 of them were detected as pA-QTLs in at least one tissue. As expected, for the vast majority of these pA-QTLs, the PAS-interrupting allele caused decreased cleavage at its p(A) site that resulted in 3’ UTR lengthening (Figure 4C). Yet, interestingly, in 12% of the cases, the effect of the PAS-interrupting allele was associated with 3’ UTR shortening, due to elevated usage of a more proximal p(A) site (Figure 4A-B). This observation argues against a scanning mechanism as the mode of action of the polyadenylation machinery. The fact that reduced efficiency of a p(A) site augments cleavage and polyadenylation at an upstream p(A) site implies a thermodynamic competition between the alternative p(A) sites.

APA potentially affects post-transcriptional gene regulation in multiple ways, including modulation of mRNA stability, translation efficiency, nuclear export, and cellular localization. However, two recent studies failed to detect a large effect of APA on transcript stability or translation efficiency [29, 30]. Yet, we sought an indication for a functional impact of the PAS SNPs on gene expression. To this goal, we intersected the 139 pA-QTLs identified by our analysis with GTEx eQTLs and found that 65 of them were also detected as eQTLs of the same gene in the same tissue. As cis-regulatory elements embedded within 3’ UTRs mainly carry destabilizing roles (e.g., miRNA binding sites, AU-rich elements (AREs)) [31], we expected that 3’ UTR lengthening effects of the PAS-interrupting alleles will be mostly linked with reduced gene expression. We indeed observed such a link in eleven out of twelve tissue (Figure 6A). On the other hand, 3’ UTR shortening effects were mostly linked with elevated gene expression (Figure 6A). Taken together, these results indicate the regulatory impact of APA on gene expression, where shorter isoforms generated by APA are generally more stable than longer ones. Nevertheless, the impact of 3’ UTR APA on transcript stability is more complicated than mere inclusion/exclusion of regulatory elements in/from the 3’ UTR, since the efficiency of mRNA targeting by such elements can be also affected by their location, as was demonstrated for miRNA target sites: sites located at the start or end of the 3’ UTRs are more efficient than those located in the middle [32]. Thus, APA can modulate the activity of miRNA target sites by changing their location relative to transcript’s 3’ end [33].

Over the last decade, GWAS studies discovered thousands of SNPs associated with common diseases and traits (The GWAS catalog already reports >70k tag SNPs [34]). Yet, the mechanism of action of most of the genetic variants identified by GWAS is currently unknown. Functional interpretation is hindered by the fact that the vast majority (>90%) of these SNPs map to noncoding regions of the genome [35]. While disruption of enhancer elements regulating gene transcription emerges as the main mode of action of risk SNPs [35], marked fractions of traits’ heritability are not accounted for by SNPs that map to transcriptional regulatory elements (e.g., putative enhancers and promoters) [36]. This indicates that other modes of action mediate the impact of noncoding genetic variants on human traits. Modulation of APA can be an important additional mechanism, and in this study, we identified 53 PAS SNPs that both significantly affect p(A) site usage (pA-QTL) and are linked to human traits (GWAS SNPs).

Our study was confined to the canonical PAS and its main variant. Examination of genetic variants affecting auxiliary APA elements is likely to identify many more such links between APA modulation, gene expression, and human traits. An analysis in this direction was very recently done by Yang et al. [37] who carried out similar pA-QTL tests using cancer RNA-seq data from The Cancer Genome Atlas (TCGA). Only 40 of the 139 pA-QTLs that we detected using GTEx data were also detected by Yang et al. using TCGA, demonstrating the complementary utility of these data resources.

Given the critical roles potentially played by APA in gene regulation and our limited understanding of how it is affected by genetic variants, our methodology and findings contribute to the initial elucidation of associations between PAS SNPs, gene expression and human phenotypes.

## Methods

### Identification of PAS SNPs

To identify PAS SNPs, we started with all 106,571 p(A) sites located in 3’ UTRs from poly(A) DB (release 3.2) [27], and all 37,611,962 annotated SNPs from GTEX v7 [28]. We considered the 53,428 SNPs located within 40-nt upstream of an annotated 3’UTR p(A) site. Examining the sequences spanning 5-nt upstream and 5-nt downstream of these SNP, we identified 2,260 instances in which one of the alleles (reference or alternative) constitute a canonic PAS sequence (AATAAA) or its main variant (ATTAAA), while the other allele interrupts this signal (we call this allele the PAS-interrupting allele - *PIA*). We also detected 45 instances in which one allele constitutes the canonic PAS, while the other allele constitutes the common variant.

### GTEx data

Aligned GTEx paired-end RNA-seq were obtained from dbGaP (release phs000424.v7.p2). We used SAMtools [38] and SRA tools to convert cram and SRA files, respectively, to BAM format. We used RNA-seq data from 12 tissues (**Tab 1**).

To determine the genotype of each PAS SNP in each RNA-seq sample, we obtained from dbGaP the processed VCF file containing the genotype of all the genomic variants analyzed by GTEx from 635 samples based on whole-genome sequencing (low-quality sites and sampled removed by GTEx analysis). We used BCF tools [39] to extract the PAS SNP genotype of each sample. eQTLs p-values and fully processed, filtered and normalized gene expression matrices were obtained from GTEx website.

### pA-QTL analysis

We associated each PAS SNP with its p(A) site (located within 40-nt downstream of it; if a PAS SNP had more than one annotated p(A) site within downstream 40-nt, the nearest p(A) site was taken). We split the 3’UTRs containing a PAS SNP into two segments: the common 3’ UTR (*cUTR*; the segment upstream of the p(A) site) and the alternative 3’ UTR (*aUTR*; the 3’ UTR segment downstream of the p(A) site). 3’ UTR coordinates were downloaded from UCSC browser based on GENCODE hg19 release v31 [40]. Overlapping 3’ UTRs of the same gene were merged using bedtools [41]. When the p(A) site of a PAS SNP was located at the edge of the 3’UTR, the entire 3’UTR was considered as the cUTR and the downstream 1,000 nt were considered as the aUTR.

We defined the pA site Usage Index (*pAUI*) to quantify the usage of each p(A) site associated with a PAS SNP in each RNA-seq sample:

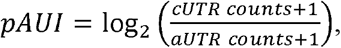

where *cUTR counts* is the fragment count in the cUTR, and *aUTR counts* is the fragment counts in the aUTR. A pseudo count of 1 was added to avoid zeroes.

We used featureCounts [42] together with a SAF file containing the coordinates of the 3’UTR segments to count the number of reads mapping to aUTRs and cUTRs in each sample. Sequenced fragments that overlapped a 3’UTR were considered in the following way: if at least one of the fragment’s reads intersected the aUTR region, the fragment was assigned to the aUTR; otherwise, the fragment was assigned to the cUTR segment. Only reads that were uniquely mapped (mapping quality of 255), aligned in proper pairs, and not marked as PCR duplicates, were counted.

We next used the pAUI levels for a pA-QTL analysis. We carried out this analysis for each tissue separately. To ensure sufficient statistical power while lowering the burden of multiple testing, for each tissue, we first filtered out PAS SNPs whose minor allele frequency (MAF) was below 0.1 among the subjects considered for that tissue. In total, over all tissue, 559 PAS SNPs (out of 2,342 PAS SNPs we detected) passed this criterion. To identify PAS SNPs that are significantly associated with the usage of their p(A) sites (that is, pA-QTL SNPs), we fitted, for each PAS SNP, a linear regression model linking its genotypes (encoded as 0, 1 and 2 according to the frequency of the PAS interrupting allele) and pAUI levels. We added as covariates to the model, the sequencing platform (Illumina HiSeq 2000 or HiSeq X), sex, and the top three genotyping principal components from GTEx analysis obtained from the GTEx website. The p-values of the genotype regression coefficients were converted to q-values to adjust for multiple testing, and a false discovery rate (FDR) threshold of 0.05 was applied to call pA-QTLs.

All tests were performed in R-3.5.1 and plots were made using ggplot2 R package.

### Association with GWAS Risk SNPs

We used Linkage disequilibrium (LD) datasets for the European population from Google Genomics. https://console.cloud.google.com/storage/browser/genomics-public-data/linkage-disequilibrium/1000-genomes-phase-3/ldCutoff0.4_window1MB/sub_pop/CEU?pli=1.

Analyzing overlaps between pA-QTLs and GWAS SNPs, we report on PAS SNPs that are in strong linkage disequilibrium (r^2^ > 0.7) with significant (p-value < 10^−5^) GWAS tag SNPs documented in the GWAS catalog (v1.0) [34].

## Supporting Information Legends

**Table S1.** 3’ UTR SNPs interrupting a canonical PAS (AATAAA) or its main variant (ATTAAA).

**Table S2.** Summary statistics for pA-QTLs detected in twelve tissues.

**Table S3.** Detailed information on the pA-QTLs and their association with eQTLs and GWAS SNPs.

